# Fitness determinants of vancomycin-resistant *Enterococcus faecium* during growth in human serum

**DOI:** 10.1101/101329

**Authors:** Xinglin Zhang, Ana M. Guzman Prieto, Vincent de Maat, Tomasz K. Prajsnar, Jumamurat R. Bayjanov, Mark de Been, Malbert R. C. Rogers, Marc J. M. Bonten, Stéphane Mesnage, Rob J. L. Willems, Willem van Schaik

## Abstract

*Enterococcus faecium* is a commensal of the human gastrointestinal tract and a frequent cause of bloodstream infections in hospitalized patients. Here, we identify genes that contribute to growth of *E. faecium* in human serum. We first sequenced the genome of *E. faecium* E745, a vancomycin-resistant clinical isolate, to completion and then compared its transcriptome during exponential growth in rich medium and in human serum by RNA-seq. This analysis revealed that 27.8% of genes on the *E. faecium* E745 genome were differentially expressed in these two conditions. A gene cluster with a role in purine biosynthesis was among the most upregulated genes in *E. faecium* E745 upon growth in serum. A high-throughput transposon sequencing (Tn-seq) approach was used to identify conditionally essential genes in *E. faecium* E745 during growth in serum. Genes involved in *de novo* nucleotide biosysnthesis (including *pyrK_2, pyrF, purD, purH*) and a gene encoding a phosphotransferase system subunit (*manY_2*) were thus identified to be contributing to *E. faecium* growth in human serum. Transposon mutants in *pyrK_2, pyrF, purD, purH* and *manY_2* were isolated from the library and their impaired growth in human serum was confirmed. In addition, the *pyrK_2* and *manY_2* mutants also exhibited significantly attenuated virulence in an intravenous zebrafish infection model. We conclude that genes involved in carbohydrate and nucleotide metabolism of *E. faecium* are essential for growth in human serum and contribute to the pathogenesis of this organism.

## Introduction

Enterococci are commensals of the gastrointestinal tract of humans and animals, but some enterococcal species, particularly *E. faecium* and *E. faecalis*, are also common causes of hospital-acquired infections in immunocompromised patients (1). While *E. faecalis* has been recognized as an important nosocomial pathogen for over a century, *E. faecium* has emerged as a prominent cause of hospital-acquired infections over the last two decades (2). Since the 1980s, *E. faecium* acquired resistance to multiple antibiotics, including β-lactams, aminoglycosides and finally, to the glycopeptide vancomycin (3). Nosocomial infections are almost exclusively caused by a specific sub-population of *E. faecium*, termed clade A-1, which has emerged from a background of human commensal and animal *E. faecium* strains (4). Strains in clade A-1 carry genetic elements that are absent from animal or human commensal isolates and which contribute to gut colonization or pathogenicity (5–9).

*E. faecium* bloodstream infections frequently develop in patients undergoing immunosuppressive therapy, and can result from translocation of strains from the intestinal tract to the bloodstream (10). In addition, the use of intravenous catheters in hospitalized patients, is another risk factor for the introduction of *E. faecium* into the bloodstream (3, 11, 12). Currently, *E. faecium* causes approximately 40% of enterococcal bacteremias. Due to the accumulation of antibiotic resistance determinants in clade A-1 strains, *E. faecium* infections are more difficult to treat than infections caused by *E. faecalis* or other enterococci (13–15).

To cause bloodstream infections, *E. faecium* needs to be able to survive and multiply in blood, but the mechanisms by which it can do so, have not yet been studied. To thrive in the bloodstream, an opportunistic pathogen has to evade host immune mechanisms and to adjust its metabolism to an environment that is relatively poor in nutrients (16).

To identify genes that are conditionally essential in bacteria, high-throughput screening methods for transposon mutant libraries have been developed and optimized for many different bacterial species (17, 18). To perform high-throughput functional genomics in ampicillin-resistant, vancomycin-susceptible clinical *E. faecium* strains, we previously developed a microarray-based transposon mutagenesis screening method which was used to identify genes involved in the development of endocarditis (7), resistance to ampicillin (19), bile (20) and disinfectants (Guzman Prieto *et al.*, submitted for publication). However, microarray-based methods for transposon mutant library screening are limited in their accuracy and can only be used in strains for which the microarray was designed. To address these limitations, several methods, including Tn-seq (21) and TraDIS (22), which are based on high-throughput sequencing of the junctions of the transposon insertion sites and genomic DNA, have been developed (23).

In this study, we set-up Tn-seq in the clinical *E. faecium* isolate E745 to identify genes that contribute to survival and growth in human serum. In addition, we determined the transcriptional response of *E. faecium* E745 in that same environment. Finally, we substantiated the role of several *E. faecium* genes that contribute to growth in serum and in virulence, in a high-throughput zebrafish infection model. Collectively, our findings show that metabolic adaptations are key to *E. faecium* growth in serum and contribute to virulence.

## Materials and Methods

### Bacterial strains, plasmids, and growth conditions

The vancomycin-resistant *E. faecium* strain E745 was used throughout this study. This strain was isolated from feces of a hospitalized patient, during a VRE outbreak in a Dutch hospital (24, 25). Unless otherwise mentioned, *E. faecium* was grown in brain heart infusion broth (BHI; Oxoid) at 37°C. The *E. coli* strains DH5*α* (Invitrogen) was grown in Luria-Bertani (LB) medium. When necessary, antibiotics were used at the following concentrations: chloramphenicol 4 µg ml^−1^ for *E. faecium* and 10 µg ml^−1^ for *E. coli*, and gentamicin 300 µg ml^−1^ for *E. faecium* and 25 µg ml^−1^ for *E. coli*. All antibiotics were obtained from Sigma-Aldrich. Growth was determined by measuring the optical density at 660 nm (OD_660_).

### Genome sequencing, assembly and bioinformatic analysis

*E. faecium* E745 was sequenced using a combination of Illumina HiSeq 100 bp paired-end sequencing, long-read sequencing using the Pacific Biosciences RS II SMRT technology and the MinION system with R7 flowcell chemistry (Oxford Nanopore Technologies). Corrected PacBio reads were assembled using the Celera assembler (version 8.1) (26) and assembled contigs were then corrected by aligning Illumina reads using BWA (version 0.7.9a), with default parameters for index creation and BWA-MEM algorithm with *-M* option for the alignment (27). This approach resulted in 15 contigs, including one contig covering the entire 2.77 Mbp chromosome. After discarding contigs with low-coverage, the remaining contigs constituted 5 circular plasmid sequences and 5 non-overlapping contigs. These 5 contigs were aligned against the NCBI Genbank database and all were found to be part of the *E. faecium* plasmid pMG1 (28). Based on this alignment the presumed order of contigs was determined and confirmed by gap-spanning PCRs and sequencing of the products. A single gap between two contigs, could not be closed by PCR. Thus, we assembled Illumina reads together with MinION 2D reads using the SPAdes assembler (version 3.0) (29), which produced a contig that closed the gap, resulting in a complete assembly of this plasmid. Sequence coverage of chromosomal and plasmid sequences was determined with SAMtools (version 0.1.18) using short read alignments to the assembly, which were generated using BWA (version 0.7.9a). SAMtools was also used to identify possible base-calling and assembly errors, by aligning short reads to assembled contigs. A base was corrected using the consensus of aligned reads (30). The corrected sequences were annotated using Prokka (version 1.10) (31). The annotated genome of *E. faecium* E745 is available from NCBI Genbank database under accession numbers CP014529 – CP014535.

### Transcriptome profiling

Approximately 3 x 10^7^ cfu of *E. faecium* E745 were inoculated into 14 ml of BHI broth and heat-inactivated serum, and grown at 37°C until log phase. Cultures were centrifuged at room temperature (15 s; 21.380 *g*), and pellets were flash frozen in liquid N_2_ prior to RNA extraction, which was performed as described previously **(19)**. The ScriptSeq Complete Kit (Bacteria) (Epicentre Biotechnologies, WI) was used for rRNA removal and strand-specific library construction. Briefly, rRNA was removed from 2.5 µg of total RNA. To generate strand specific RNA-seq data, approximately 100 ng of rRNA-depleted RNA was fragmented and reverse transcribed using random primers containing a 5′ tagging sequence, followed by 3′ end tagging with a terminal-tagging oligo to yield di-tagged, single-stranded cDNA. Following magnetic-bead based purification, the di-tagged cDNA was amplified by PCR (15 cycles) using ScriptSeq Index PCR Primers (Epicentre Biotechnologies, WI). Amplified RNA-seq libraries were purified using AMPure XP System (Beckman Coulter) and sequenced by a 100 bp paired end reads sequencing run using the Illumina HiSeq 2500 platform (University of Edinburgh, United Kingdom). Data analysis was performed using Rockhopper **(32)** using the default settings for strand specific analysis.

### Confirmation of RNA-seq data by quantitative real-time RT-PCR (qRT-PCR)

Total RNA isolated as described previously was used to confirm the transcriptome analysis by qRT-PCR. cDNA was synthesized as described above and qRT-PCR on these cDNAs was performed using the Maxima SYBR Green/ROX qPCR Master Mix (Thermo Scientific, Breda, The Netherlands) and a StepOnePlus instrument (Life Technologies). The expression of *tufA* was used as a housekeeping control. Ct values were calculated using the StepOne analysis software v2.2. Transcript levels, relative to *tufA*, of the assayed genes were calculated using REST 2009 V2.0.13 (Qiagen, Venlo, The Netherlands). This experiment was performed with three biological replicates.

### Generation of *mariner* transposon mutant library in *E. faecium*

To create a transposon mutant library in *E. faecium* E745 suitable for Tn-seq, the *mariner* transposon cassette (carrying a gentamicin resistance gene) in the transposon delivery plasmid pZXL5 (19) was adapted as follows. The transposon from pZXL5 was amplified by PCR using the set of primers: pZXL5_MmeI_SacII_Fw and pZXL5_MmeI_SacII_Rv (primer sequences listed in Supplementary table 7). These primers introduced MmeI restriction sites in the inverted repeats on both sides of the transposon. The modified transposon delivery vector, termed pGPA1, was generated by the digestion of pZXL5 with SacII, followed by the insertion of the SacII-digested *mariner* transposon that contained MmeI restriction sites at its extreme ends. pGPA1 was electroporated into *E. faecium* E745 and the transposon mutant library was generated by selecting for gentamicin-resistant transposon mutants as described previously (19).

### Tn-seq analysis of conditionally essential genes in *E. faecium* E745

The transposon mutant library created in E745 was prepared for Tn-seq analysis, similar to previously described procedures **(33)**. To identify genes that are essential for the viability of *E. faecium* in BHI, we used ten experimental replicates of the mutant library. Aliquots (20 μl) of the transposon mutant library, containing approximately 10^7^ cfu, were used to inoculate 20 ml BHI broth and grown overnight at 37°C. Subsequently, 1 ml aliquots of the cultures were spun down (15 s, 21.380 *g*) and used for the extraction of genomic DNA (Wizard genomic DNA purification kit, Promega Benelux). 2 μg of the extracted DNA was digested for 4 hr at 37°C using 10U MmeI (New England Biolabs) and immediately dephosphorylated with 1U of calf intestine alkaline phosphatase (Invitrogen) during 30 min at 50°C. DNA was isolated using phenol-chloroform extraction and subsequently precipitated using ethanol. The DNA pellets were then dissolved in 20 μl water. The samples were barcoded and prepared for Tn-seq sequencing as described previously **(33)**. The sequence reads from all ten experimental replicates were mapped to the genome, and the mapped read-counts were then tallied for the analysis of the essentiality of the genes in the *E. faecium* E745 genome (further described below).

To identify genes that are required for growth in human serum, 20 μl aliquots of the frozen mutant library in E745 were inoculated in BHI broth and grown overnight as described above. Subsequently, bacterial cells were washed with physiological saline solution. Approximately 3x10^7^ cfu were inoculated into 14 ml BHI broth, and approximately 3x10^6^ cfu were inoculated into 14 ml human serum obtained from Sigma (Cat. No. H4522; Sterile filtered type-AB human serum) or heat-inactivated human serum (the same, after incubation for 30 min at 56°C). The different inoculum-sized were used in order for a similar number of divisions to occur during the experiment. Cells were incubated at 37°C for 24 hours without shaking and then further processed for Tn-seq (33). This experiment was performed in triplicate.

Tn-seq samples were sequenced (50 nt, single-end) on one lane of a Illumina Hiseq 2500 (Baseclear, Leiden, the Netherlands and Sequencing facility University Medical Center, Utrecht, The Netherlands), generating an average of 15 million high quality reads per sample.

### Tn-seq data analysis

Raw Illumina sequence reads from Illumina sequencing were split, based on their barcode, using the Galaxy platform (34), and 16-nucleotide fragments of each read that corresponded to E745 sequences, were mapped to the E745 genome using Bowtie 2 (35). The results of the alignment were sorted and counted by IGV (36) using a 25-nucleotide window size and then summed over the gene. Read mapping to the final 10% of a gene were discarded as these insertions may not inactivate gene function. Read counts per gene were then normalized to the total number of reads that mapped to the genome in each replicate, by calculating the normalized read-count RPKM (Reads Per Kilobases per Million input reads) via the following formula: RPKM = (number of reads mapped to a gene x 10^6^) / (total mapped input reads in the sample x gene length in kbp). Statistical analysis of the RPKM-values between the experimental conditions was performed using Cyber-T (37). Genes were determined to be significantly contributing to growth in human serum when the Benjamini-Hochberg corrected *P-*value was <0.05 and the difference in abundance of the transposon mutant during growth in BHI and serum was >2. To determine the essentiality of *E.faecium* genes during growth in the rich medium BHI, the normalized read-counts of the ten replicates in BHI were used as data input for the EL-ARTIST analysis, as described in the user manual of the ARTIST pipeline (38).

### Isolation of mutants from the transposon mutant library pool

To recover a targeted transposon mutant from the complete mutant pool, a PCR-based screening strategy was developed (Supplementary figure 3). 40 µl of the transposon mutant library was inoculated into 40 ml of BHI broth with gentamicin and grown overnight at 37ºC with shaking (200 rpm). The overnight culture, containing approximately 10^9^ cfu/ml, was then diluted to approximately 20 cfu/ml in 500 ml of BHI with gentamicin and kept on ice. Subsequently, 200 μl aliquots were transferred to wells of sterile 96 wells plates (n = 12, Corning Inc.). After overnight incubation at 37ºC without shaking, aliquots (15 μl) of each one of the 96 wells, were further pooled into a single new 96 well plate, as described in Supplementary figure 3.

PCRs were performed on the final plate in which the transposon mutants were pooled, to check for the presence of the Tn-mutants of interest, using the primer ftp_tn_both_ends_MmeI, which is complementary to the repeats flanking the transposon sequence, in combination with a gene-specific primer (Supplementary table 7). When a PCR was found to be positive in one of the wells of this plate, the location of the Tn-mutant was tracked backwards to the wells containing approximately 4 independent transposon mutants, by performing PCRs mapping the presence of the transposon mutant in each step. Cells from the final positive well were plated onto BHI with gentamicin and colony PCR was performed to identify the desired transposon mutant.

### Growth of *E. faecium* E745 and individual mutants in human serum

Wild-type E745 and the mutant strains were grown overnight at 37°C in BHI broth. Subsequently, bacterial cells were washed with physiological saline and approximately 3 x 10^5^ cfu were inoculated into 1.4 ml BHI broth or heat-inactivated serum. Cells were grown in 1.5 ml tubes (Eppendorf) in triplicate for each condition and incubated at 37°C for 24 hours without shaking. Bacterial growth was determined by assessing viable counts, for which the cultures were serially diluted using physiological saline solution and plated onto BHI agar followed by overnight incubation at 37°C.

### Intravenous infection of zebrafish embryos

London wild-type (LWT) inbred zebrafish embryos, provided by the aquarium staff of The Bateson Center (University of Sheffield), were used for infection experiments. The parental E745 strain and its *pyrK_2* and *manY_2* transposon mutants were grown in BHI broth until they reached an optical density at 600 nm of approximately 0.5 and were then harvested by centrifugation (5,500 *g*, 10 min). Bacteria were microinjected into the circulation of dechorionated zebrafish embryos at 30 hours post fertilization, as previously described **(39)**. Briefly, anesthetized embryos were embedded in 3% (w/v) methylcellulose and injected individually with approximately 1.2 x 10^4^ cfu using microcapillary pipettes. For each strain, 29 to 32 infected embryos were observed for survival up to 90 hours post infection (hpi). This experiment was performed in triplicate.

### Data availability

Sequence reads generated in this study have been made available at the European Nucleotide Archive under accession number PRJEB19025.

## Results

### The complete genome sequence of *E. faecium* E745

In this study, we implemented RNA-seq and Tn-seq analyses in *E. faecium* strain E745, an ampicillin-and vancomycin-resistant clinical isolate that was isolated from faeces of a hospitalized patient during an outbreak of VRE in the nephrology ward of a Dutch hospital in 2000 (24, 25). To allow the application of RNA-seq and Tn-seq in *E. faecium* E745, we first determined the complete genome sequence of this strain through a combination of short-read Illumina sequencing and long-read sequencing on the RSII Pacific Biosciences and Oxford NanoPore’s MinION systems. This resulted in a circular chromosomal sequence of 2,765,010 nt and 6 plasmids, with sizes ranging between 9.3 kbp and 223.7 kbp (Supplementary table 1). Together, the chromosome and plasmids have 3,095 predicted coding sequences.

### Transcriptome of *E. faecium* E745 during growth in rich medium and in human serum

The transcriptional profile of E745 was determined using RNA-seq during growth in rich medium (BHI) and in heat-inactivated human serum. A total of 99.9 million (15.6 - 17.6 million per sample) 100 bp paired-end reads were successfully aligned to the genome, allowing the quantification of rare transcripts (Fig. 1). A total of 3217 transcription units were identified, including 651 predicted multi-gene operons, of which the largest contains 22 genes (Fig. 1A and Supplementary table 2).

**Figure 1:**
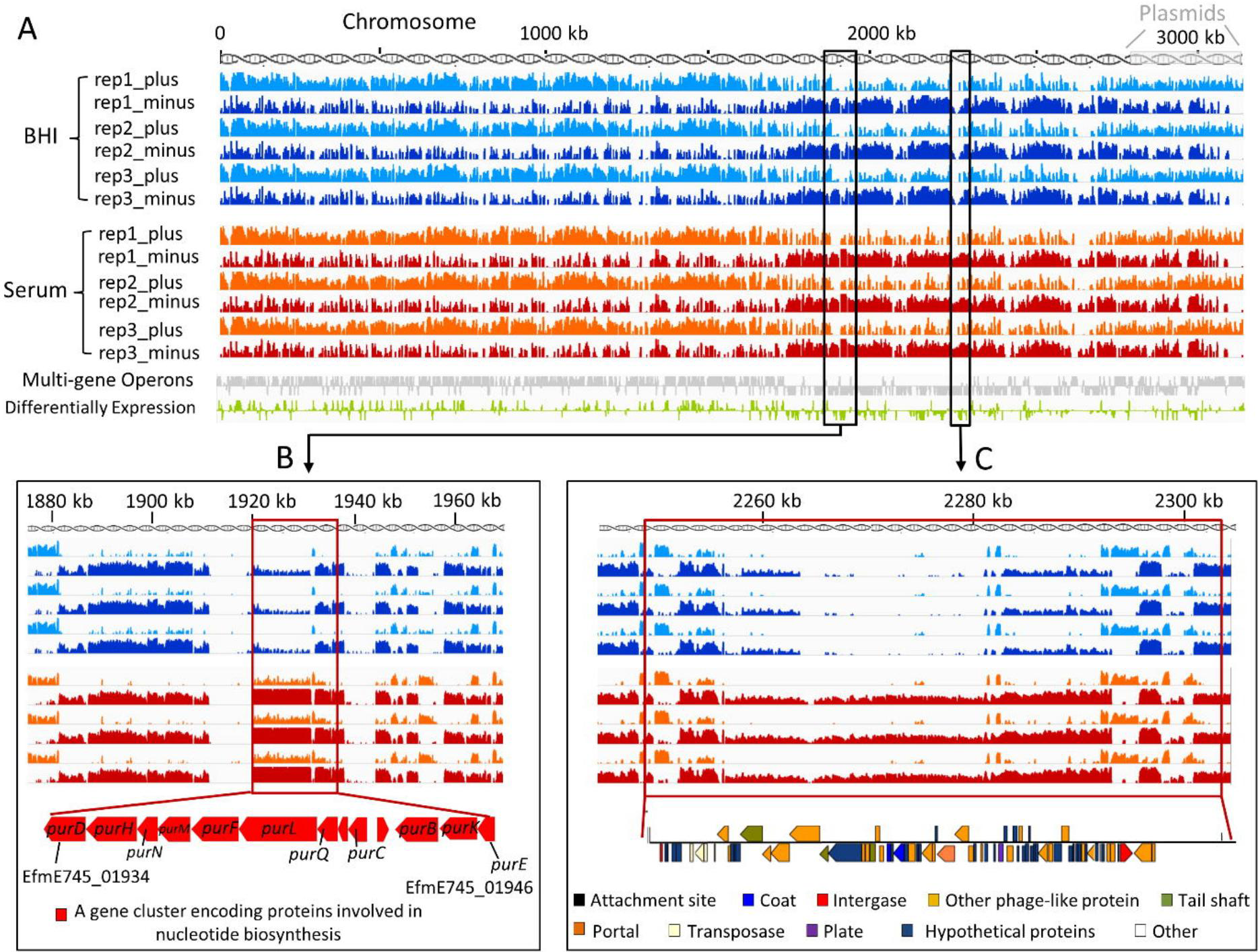
Coverage plots of RNA-seq data aligning to chromosome and plasmid DNA. The *y* axis of each track indicates reads coverage and is represented on a log scale, ranging from 0 to 10000. The *x* axis represents the genomic location. Light blue (BHI) or orange (serum) tracks correspond to sequencing reads aligned to the plus strand of the replicon, and dark blue (BHI) or dark red (serum) tracks correspond to sequencing reads aligned to the minus strand of the replicon. The grey track corresponds to multi-gene operons. The green track corresponds to differentially expressed genes (BHI vs serum), with the height of the green bars indicative of differential expression. In panels B and C, two serum-induced regions are shown, i.e. a gene cluster involved in nucleotide biosynthesis (panel B) and a prophage (panel C). The RNA-seq experiments were performed using three biological replicates.

A comparative analysis of E745 during growth in BHI and in human serum, showed that 860 genes exhibited significantly (*q*<0.001 and a fold change in expression of >2 or <0.5 between cultures grown in BHI versus heat-inactivated serum) different expression between these conditions (Supplementary table 3). The large number (27.8% of genes on the *E. faecium* E745 genome) of differentially expressed genes, indicates that growth in human serum leads to a dramatic reprogramming of global *E. faecium* gene expression, involving both chromosomal and plasmid-located genes. Among the genes with the highest difference in expression between growth in serum and in rich medium, we identified a gene cluster with a role in purine biosynthesis (Fig. 1B). In addition, we found a 58.4 kbp prophage-like gene cluster that exhibited higher expression in E745 during growth in serum (Fig. 1C).

To confirm the RNA-seq analysis, we independently determined expression levels of eight genes during growth in serum versus growth in BHI by qPCR (Supplementary Fig. 1). RNA-seq and qPCR data were highly concordant (r^2^ = 0.98).

### Construction and analysis of a transposon mutant library in *E. faecium* E745

A *mariner*-based transposon mutant library was generated in *E. faecium* E745 and Tn-seq (21) was performed on ten replicate transposon mutant libraries, that were grown overnight in BHI at 37°C, resulting in an average of 15 million Tn-seq reads for each library. To analyze the Tn-seq data, we divided the E745 genome in 25-nt windows. Of a total of 110,601 windows, 49,984 (45%) 25-nt windows contained one or more sequence reads. No positional bias was observed in the transposon insertion sites in the chromosome and plasmids of *E.faecium* E745 (Supplementary Fig. 2A). The genome-wide coverage of this transposon mutant library allowed the identification of genes that are conditionally essential in *E.faecium*. A total of 455 chromosomal genes were determined to be essential for growth of *E.faecium* E745 in BHI (Supplementary Table 4). An example of a gene identified as essential is shown in Supplementary Fig. 2B. Among the genes that were essential for growth of *E.faecium* E745 in BHI, 87% were present in all genome sequences of a set of 74 previously sequenced *E. faecium* strains (4) that represented the genetic and ecological diversity of the species. An additional 7% of the essential genes were present in 73 out of 74 *E. faecium* genomes (data not shown). The conserved presence of these essential genes among diverse *E.faecium* strains is in line with their crucial role in *E. faecium* viability. The *E. faecium* E745 transposon mutant library was then used to identify genes that were specifically required for growth of *E. faecium* in serum.

### *E. faecium* E745 genes required for growth in human serum

In order to identify genes that contribute to growth of *E. faecium* E745 in human serum, we performed Tn-seq on cultures of the *E. faecium* E745 transposon mutant libraries upon growth in rich medium (BHI) and in human serum. The human serum was either used natively or heat-treated. The serum showed effective complement activity in both classical and alternative pathways, as determined by hemolytic assays (40), while this activity was abolished by heat-inactivation (data not shown). Minor differences were observed among conditionally essential genes between the experiments performed in native human serum or heat-inactivated serum (Supplementary Table 5) and the following results correspond to the experiments obtained with heat-inactivated serum. This condition was chosen because it may be a more reproducible *in vitro* environment, particularly since the interaction between the complement system and Gram-positive bacteria remains to be fully elucidated (41, 42).

We identified 37 genes that significantly contributed to growth of E745 in human serum (Fig. 2 and Supplementary table 6): twenty-nine genes were located on the chromosome and eight genes were present on plasmids (six genes on pE745-5, two genes on plasmid pE745-6). The relatively large number of genes identified indicates that growth of *E. faecium* in human serum is a multi-factorial process. The genes that conferred the most pronounced effect on growth of *E. faecium* in serum included genes that are part of a phosphotransferase systems (PTS) involved in carbohydrate uptake (*manZ_3, manY_2*, *ptsL*), a putative transcriptional regulator (*algB*) and genes involved in the biosynthesis of purine and pyrimidine nucleotides (*guaB*, *purA*, *pyrF*, *pyrK_2*, *purD*, *purH*, *purL*, *purQ*, *purC*) (Fig. 2). Notably, the *purD, purH*, and *purL* genes were found to exhibit higher expression upon growth in human serum in the RNA-seq analysis (Fig. 1). Nine genes were identified as negatively contributing to growth in serum, i.e. the transposon mutants in these genes were significantly enriched upon growth in serum. The effects of these mutations were relatively limited, compared to the major effects observed in the transposon mutants discussed above (Supplementary Table 6), but it is notable that five (*clsA_1*, *ddcP, ldt*_*fm*_*, mgs*, and *lytA_2*) of these genes have roles in cell wall and cytoplasmic membrane biosynthesis.

**Figure 2:**
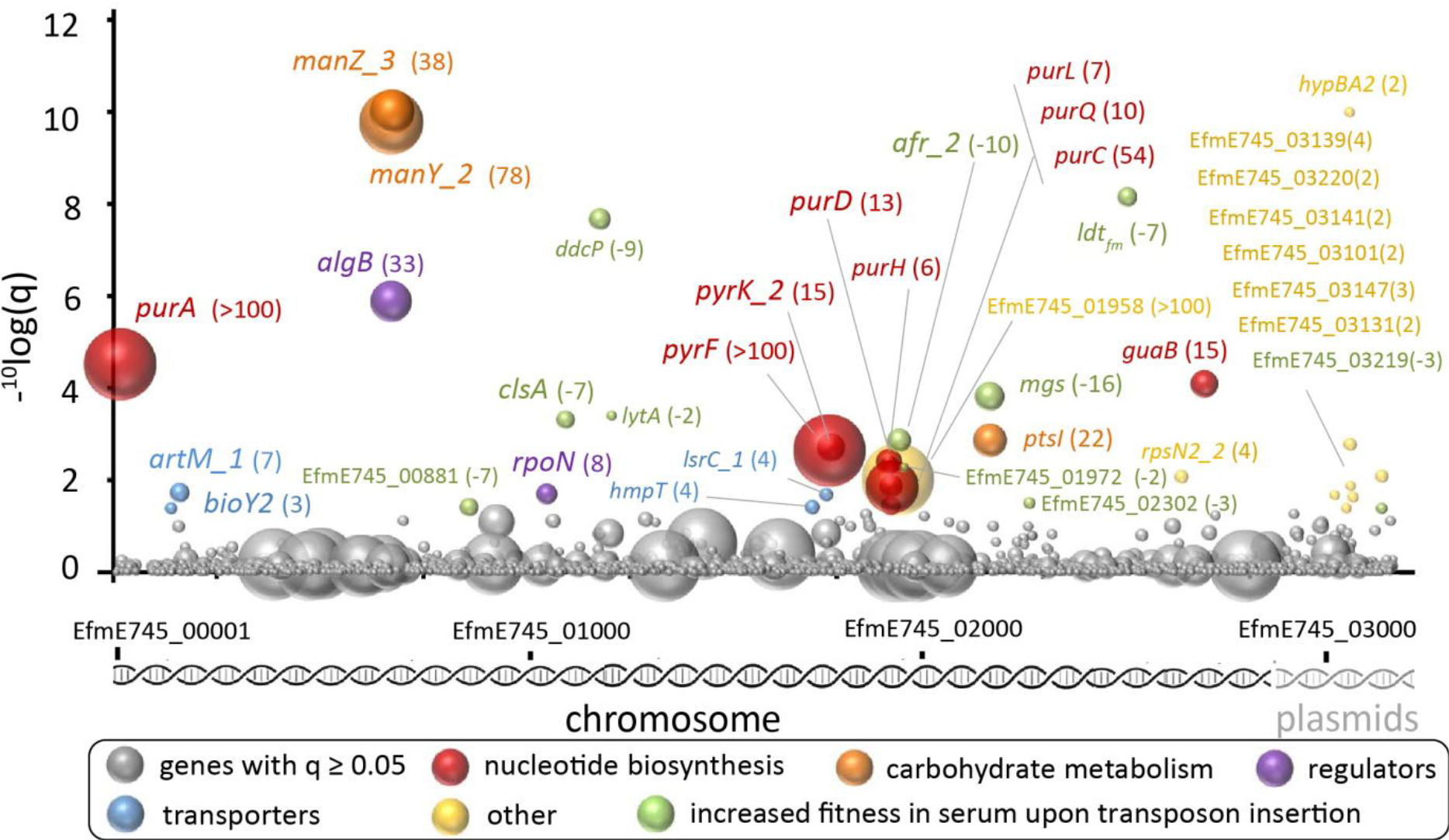
Tn-seq analysis for *E. feacium*genes required for growth in human serum. Bubbles represent genes, and bubble size corresponds to the fold-changes (for visual reasons, a 100-fold change in transposon mutant abundance is set as a maximum) derived from the read-count ratio of libraries grown in BHI to libraries grown in human serum. On the *x* axis genes are shown in order of their genomic location and the chromosome and plasmids are indicated. The outcome of statistical analysis of the Tn-seq data is indicated on the *y* axis. Genes with a significant change (*q* < 0.05) in fitness in serum versus BHI are grouped by function and are labelled with different colors, and the name or locus tag and the change in abundance between the control condition and growth in serum is indicated next to the bubbles. Negative values indicate that mutants in these genes outgrow other mutants in serum, suggesting that these mutants, compared to the wild-type E745, have a higher fitness in serum.

### *E. faecium* genes involved in nucleotide biosynthesis or carbohydrate metabolism contribute to growth in human serum

We developed a PCR-based method (Supplementary Fig. 3) to selectively isolate five transposon mutants (in the purine metabolism genes *purD* and *purH*, the pyrimidine metabolism genes *pyrF* and *pyrK_2* and the PTS gene *manY_2*) from the transposon library. Growth in rich medium of these transposon insertion mutants was equal to the parental strain. However, all mutants were significantly impaired in their growth in human serum (Fig. 3A), confirming the results of the Tn-seq experiments.

**Figure 3.**
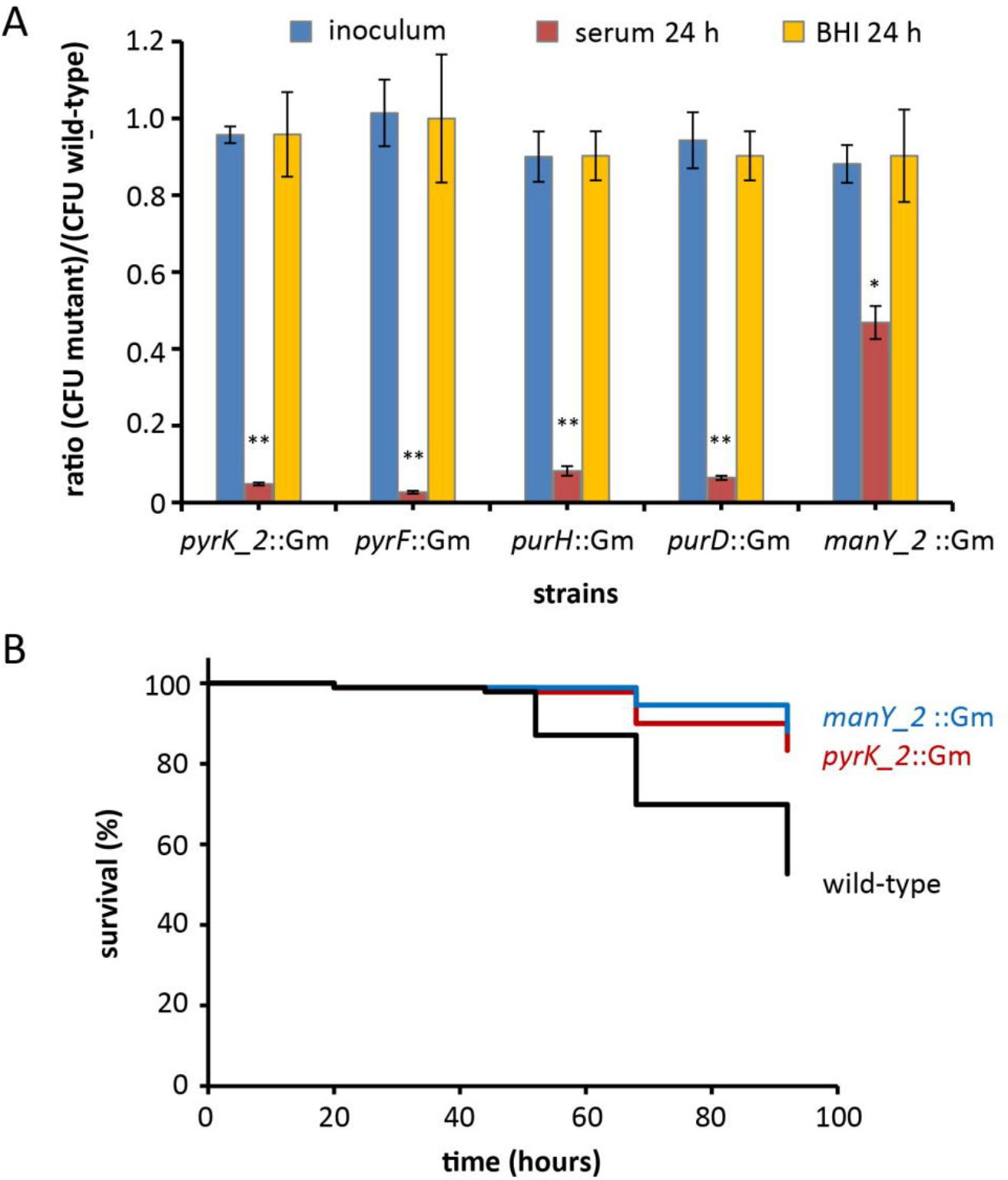
*E. faecium* transposon mutants with a growth defect in human serum and an attenuated phenotype in a zebrafish model. (A) Ratios of the viable counts of five mutantscompared to wild-type *E. faecium* before (blue bars) and after 24 h of growth in human serum (red bars) or BHI (yellow bars). The viable counts of wild-type *E. faecium* E745 were (3.52 ± 0.07) x 10^5^/ml in the inocula, (2.92 ± 0.14) x 10^8^/ml after 24 h of growth in serum and (1.20 ± 0.20) x 10^9^/ml after 24 h of growth in BHI, respectively. Error bars represent the standard deviation of the mean of three independent experiments. Asterisks represent significant difference (*: *p* < 10^-3^,**: *p* < 10^-5^ as determined by a two-tailed Student’s *t*-test) between the mutant strains and wild-type. (B) Kaplan-Meier survival curves of zebrafish embryos upon infection with *E. faecium*. The virulence of *E. faecium* mutants upon intravenous infection of zebrafish embryos was determined upon injection with 1.2x10^4^ cfu of the *manY_2*: Gm and *pyrK*: Gm transposon mutants and wild-type *E. faecium* E745. The experiment was performed three times and the mutants were significantly different (*p*<0.01) from the wild-type in each experiment as determined by the Log-rank (Mantel-Cox) test with Bonferroni correction for multiple comparisons. This figure represents the combined results of the three replicates for *E. faecium* E745 (*n =* 93 zebrafish embryos), *manY_2*: Gm (*n =* 92) and *pyrK*: Gm (*n =* 90).

### *E. faecium pyrK_2* and *manY_2* contribute to intravenous infection of zebrafish

Next, we investigated whether the transposon insertion mutants in the *manY_2* and *pyrK_2* genes were attenuated *in vivo* (Fig. 3B). The mutants in these genes were selected because they represent the mutants in nucleotide and carbohydrate metabolism genes that were previously shown to contribute to the growth of *E. faecium* in human serum. As a model for intravenous infection, we used a recently described model in which *E. faecium* was injected into the circulation of zebrafish embryos to mimic systemic infections (43). We showed that both the *manY_2* and the *pyrK_2* mutant were significantly less virulent than the parental strain. The overall survival at 92 hours post infection (hpi) of zebrafish embryos infected with WT strain was 53%, compared to 88% and 83% respectively, for zebrafish embryos that were infected with the transposon insertion mutants in *manY_2* and *pyrK_2*.

## Discussion

*E. faecium* can contaminate the skin and from there colonize indwelling devices such as intravenous catheters, or it can translocate from the gastrointestinal tract in immunosuppressed patients, leading to the development of bacteremia and endocarditis. *E.faecium* infections are often difficult to treat, due to the multi-drug resistant character of the strains causing nosocomial infections (3, 4). However, the bloodstream poses challenges for the proliferation and survival of *E. faecium*, including a scarcity of nutrients.

In the present study we sequenced the complete genome of a vancomycin-resistant *E. faecium* strain, and identified *E. faecium* genes that were essential for growth in human serum. A total of 37 genes were found to be required for fitness of *E. faecium* E745 in serum, among which genes with roles in carbohydrate uptake and nucleotide biosynthesis. Previously, fitness determinants for growth in human serum have been identified through large-scale screening of mutant libraries in both a Gram-negative (*Escherichia coli*) and a Gram-positive (*Streptococcus pyogenes*) pathogen (44, 45). Notably, these studies have also identified the ability for *de novo* synthesis of purines and pyrimidines as a crucial factor for growth in serum. In addition, in diverse bacteria (including *Burkholderia cepacia, Pasteurella multocida*, *Acinetobacter baumannii*, *Salmonella enterica* serovar Typhimurium, *Bacillus anthracis*, and *Streptococcus pneumoniae*), the ability to synthesize nucleotides contributes importantly to virulence (46–51). The data presented here indicate that *de novo* biosynthesis of nucleotides is also required for *E. faecium* growth in serum and virulence. The nucleotide biosynthesis pathway of *E. faecium* may be a promising target for the development of novel antimicrobials for the treatment of *E. faecium* bloodstream infections. Indeed, compounds that target guanine riboswitches, thereby inhibiting nucleotide biosynthesis, have already shown their efficacy in a *Staphylococcus aureus* infection model (52).

Three genes (*ptsL*, *manY_2* and *manZ_3*) encoding subunits of PTSs were found to contribute to growth in serum in our Tn-seq experiments. Previously, PTSs have been associated with gut colonization (5) and endocarditis (7) in another clinical *E. faecium* strain. The PTSs identified in this study are different from these previously characterized systems, suggesting that the remarkably large number of genes encoding for PTSs in a typical *E. faecium* genome (4) provide metabolic flexibility for growth in a wide variety of environments.

It is notable that among the nine genes that exhibited increased fitness upon inactivation by transposon insertion, five genes are predicted to have a role in cell wall or cytoplasmic membrane biosynthesis. The protein encoded by *ddcP* was previously characterized as a low-molecular-weight penicillin binding protein with D-alanyl-D-alanine carboxypeptidase activity (19), while *ldt*_*fm*_ acts as a peptidoglycan L, D transpeptidase (53). The predicted α-monoglucosyldiacylglycerol synthase gene *mgs* is orthologous (73% amino acid identity) to *bgsB* in *E. faecalis*, which is required for the biosynthesis of membrane glycolipids (54). The *clsA_1* gene is predicted to be responsible for the synthesis of cardiolipin (bisphosphatidylglycerol) and its inactivation may modulate the physical properties of the cytoplasmic membrane (55). Finally, *lytA_2* is predicted to encode an autolysin, which may be involved in the turnover of peptidoglycan in the cell wall (56). The transposon mutants in these genes were not further characterized in this study, but our findings suggest that genes involved in cell wall or cytoplasmic membrane remodeling may confer subtle fitness defects to *E. faecium* when growing in serum.

Our RNA-seq-based transcriptional profiling of *E. faecium* E745 during mid-exponential growth in serum showed pervasive changes in gene expression compared to exponential growth in rich medium. The purine metabolism genes *purL*, *purH*, *purD*, which were found to be required for growth in serum in our Tn-seq experiments, were among those that were significantly upregulated during growth in serum compared to growth in rich medium. Notably, a single prophage was expressed at higher levels during growth in serum than in rich medium. The abundance of prophage elements in the genome of *E. faecium* has been noted before (4, 57, 58). Interestingly, in the related bacterium *Enterococcus faecalis* prophages encode platelet-binding proteins (59) and may have a role in intestinal colonization (60). The contribution of *E. faecium* prophages to traits that are important for colonization and infection may provide important insights into the success of *E. faecium* as a nosocomial pathogen.

Collectively our data indicates that nucleotide biosynthesis and carbohydrate metabolism are critical metabolic pathways for the proliferation and survival of *E. faecium* in the bloodstream. The proteins encoded by the genes required for growth in human serum that were identified in this study, could serve as candidates for the development of novel anti-infectives for the treatment of bloodstream-infections by multi-drug resistant *E. faecium*.

## Acknowledgements

This work was supported by the European Union Seventh Framework Programme (FP7-HEALTH-2011-single-stage) “Evolution and Transfer of Antibiotic Resistance” (EvoTAR) under grant agreement number 282004 and by an NWO-VENI grant (916.86.044) and NWO-VIDI grant (917.13.357) to W.v. S.

## Author contributions

X. Z, R. J. L. W. and W.v. S. designed the study. X. Z., A. M. G. P., T. K. P., M. B., J. R. B., M. R. and S. M. performed experiments. All authors contributed to data interpretation. The manuscript was written by Z. X., A. M. G. P., M. J. M. B., R. J. L. W. and W.v. S.

## References

1. Dupont, H., Friggeri, A., Touzeau, J., Airapetian, N., Tinturier, F., Lobjoie, E., Lorne, E., Hijazi, M., Régimbeau,J.-M. and Mahjoub, Y. (2011) Enterococci increase the morbidity and mortality associated with severe intra-abdominal infections in elderly patients hospitalized in the intensive care unit. J. Antimicrob. Chemother., 66, 2379–2385.

2. Guzman Prieto, A.M., van Schaik, W., Rogers, M.R.C., Coque, T.M., Baquero, F., Corander, J. and Willems, R.J.L. (2016) Global emergence and dissemination of enterococci as nosocomial pathogens: attack of the clones? Front. Microbiol., 7, 788.

3. Arias, C.A. and Murray, B.E. (2012) The rise of the Enterococcus: beyond vancomycin resistance. Nat. Rev. Microbiol., 10, 266–278.

4. Lebreton, F., van Schaik, W., Manson McGuire, A., Godfrey, P., Griggs, A., Mazumdar, V., Corander, J., Cheng, L., Saif, S., Young, S., et al. (2013) Emergence of epidemic multidrug-resistant Enterococcus faecium from animal and commensal strains. mBio, 4, e00534–13.

5. Zhang, X., Top, J., de Been, M., Bierschenk, D., Rogers, M., Leendertse, M., Bonten, M.J.M., van der Poll, T., Willems, R.J.L. and van Schaik, W. (2013) Identification of a genetic determinant in clinical Enterococcus faecium strains that contributes to intestinal colonization during antibiotic treatment. J. Infect. Dis., 207, 1780–1786.

6. Heikens, E., Singh, K.V., Jacques-Palaz,K.D., van Luit-Asbroek,M., Oostdijk, E.A.N., Bonten, M.J.M., Murray, B.E. and Willems, R.J.L. (2011) Contribution of the enterococcal surface protein Esp to pathogenesis of Enterococcus faecium endocarditis. Microbes Infect., 13, 1185–1190.

7. Paganelli, F.L., Huebner, J., Singh, K.V., Zhang, X., van Schaik, W., Wobser, D., Braat, J.C., Murray, B.E., Bonten, M.J.M., Willems, R.J.L., et al. (2016) Genome-wide screening identifies phosphotransferase system permease BepA to be Involved in Enterococcus faecium endocarditis and biofilm formation. J. Infect. Dis., 214, 189–195.

8. Sillanpää,J., Prakash, V.P., Nallapareddy, S.R. and Murray, B.E. (2009) Distribution of genes encoding MSCRAMMs and pili in clinical and natural populations of Enterococcus faecium. J. Clin. Microbiol., 47, 896–901.

9. Montealegre, M.C., Singh, K.V., Somarajan, S.R., Yadav, P., Chang, C., Spencer, R., Sillanpää,J., Ton-That,H. and Murray, B.E. (2016) Role of the Emp pilus subunits of Enterococcus faecium in biofilm formation, adherence to host extracellular matrix components, and experimental infection. Infect. Immun., 84, 1491–1500.

10. Kamboj, M., Blair, R., Bell, N., Sun, J., Eagan, J. and Sepkowitz, K. (2014) What is the source of bloodstream infection due to vancomycin-resistant enterococci in persons with mucosal barrier injury? Infect. Control Hosp. Epidemiol., 35, 99–101.

11. Bouza, E., Kestler, M., Beca, T., Mariscal, G., Rodríguez-Créixems,M., Bermejo, J., Fernández-Cruz,A., Fernández-Avilés,F., Muñoz,P. and Grupo de Apoyo al Manejo de la Endocarditis (2015) The NOVA score: a proposal to reduce the need for transesophageal echocardiography in patients with enterococcal bacteremia. Clin. Infect. Dis., 60, 528–535.

12. Arias, C.A. and Murray, B.E. (2008) Emergence and management of drug-resistant enterococcal infections. Expert Rev. Anti Infect. Ther., 6, 637–655.

13. Coombs, G.W., Pearson, J.C., Daly, D.A., Le, T.T., Robinson, J.O., Gottlieb, T., Howden, B.P., Johnson, P.D.R., Bennett, C.M., Stinear, T.P., et al. (2014) Australian enterococcal sepsis outcome programme annual report, 2013. Commun. Dis. Intell. Q. Rep., 38, E320–326.

14. de Kraker, M.E.A., Jarlier, V., Monen, J.C.M., Heuer, O.E., van de Sande, N. and Grundmann, H. (2013) The changing epidemiology of bacteraemias in Europe: trends from the European Antimicrobial Resistance Surveillance System. Clin. Microbiol. Infect., 19, 860–868.

15. Hidron, A.I., Edwards, J.R., Patel, J., Horan, T.C., Sievert, D.M., Pollock, D.A., Fridkin, S.K., National Healthcare Safety Network Team and Participating National Healthcare Safety Network Facilities (2008) NHSN annual update: antimicrobial-resistant pathogens associated with healthcare-associated infections: annual summary of data reported to the National Healthcare Safety Network at the Centers for Disease Control and Prevention, 2006-2007. Infect. Control Hosp. Epidemiol., 29, 996–1011.

16. Krebs, H.A. (1950) Chemical composition of blood plasma and serum. Annu. Rev. Biochem., 19, 409–430.

17. van Opijnen, T. and Camilli, A. (2013) Transposon insertion sequencing: a new tool for systems-level analysis of microorganisms. Nat. Rev. Microbiol., 11, 435–442.

18. Barquist, L., Boinett, C.J. and Cain, A.K. (2013) Approaches to querying bacterial genomes with transposon-insertion sequencing. RNA Biol., 10, 1161–1169.

19. Zhang, X., Paganelli, F.L., Bierschenk, D., Kuipers, A., Bonten, M.J.M., Willems, R.J.L. and van Schaik, W. (2012) Genome-wide identification of ampicillin resistance determinants in Enterococcus faecium. PLoS Genet., 8, e1002804.

20. Zhang, X., Bierschenk, D., Top, J., Anastasiou, I., Bonten, M.J., Willems, R.J. and van Schaik, W. (2013) Functional genomic analysis of bile salt resistance in Enterococcus faecium. BMC Genomics, 14, 299.

21. van Opijnen, T., Bodi, K.L. and Camilli, A. (2009) Tn-seq: high-throughput parallel sequencing for fitness and genetic interaction studies in microorganisms. Nat. Methods, 6, 767–772.

22. Langridge, G.C., Phan, M.-D., Turner, D.J., Perkins, T.T., Parts, L., Haase, J., Charles, I., Maskell, D.J., Peters, S.E., Dougan, G., et al. (2009) Simultaneous assay of every Salmonella Typhi gene using one million transposon mutants. Genome Res., 19, 2308–2316.

23. van Opijnen, T., Lazinski, D.W. and Camilli, A. (2015) Genome-wide fitness and genetic interactions determined by Tn-seq, a high-throughput massively parallel sequencing method for microorganisms., Current Protocols in Microbiology. John Wiley & Sons, Inc, Hoboken, NJ, USA, p.1E.3.1-1E.3.24.

24. Mascini, E.M., Troelstra, A., Beitsma, M., Blok, H.E.M., Jalink, K.P., Hopmans, T.E.M., Fluit, A.C., Hene, R.J., Willems, R.J.L., Verhoef, J., et al. (2006) Genotyping and preemptive isolation to control an outbreak of vancomycin-resistant Enterococcus faecium. Clin. Infect. Dis., 42, 739–746.

25. Leavis, H.L., Willems, R.J.L., van Wamel, W.J.B., Schuren, F.H., Caspers, M.P.M. and Bonten, M.J.M. (2007) Insertion sequence-driven diversification creates a globally dispersed emerging multiresistant subspecies of E. faecium. PLoS Pathog., 3, e7.

26. Goldberg, S.M.D., Johnson, J., Busam, D., Feldblyum, T., Ferriera, S., Friedman, R., Halpern, A., Khouri, H., Kravitz, S.A., Lauro, F.M., et al. (2006) A Sanger/pyrosequencing hybrid approach for the generation of high-quality draft assemblies of marine microbial genomes. Proc. Natl. Acad. Sci. U. S. A., 103, 11240–11245.

27. Li, H. and Durbin, R. (2009) Fast and accurate short read alignment with Burrows-Wheeler transform. Bioinformatics, 25, 1754–1760.

28. Tanimoto, K. and Ike, Y. (2008) Complete nucleotide sequencing and analysis of the 65-kb highly conjugative Enterococcus faecium plasmid pMG1: identification of the transfer-related region and the minimum region required for replication. FEMS Microbiol. Lett., 288, 186–195.

29. Bankevich, A., Nurk, S., Antipov, D., Gurevich, A.A., Dvorkin, M., Kulikov, A.S., Lesin, V.M., Nikolenko, S.I., Pham, S., Prjibelski, A.D., et al. (2012) SPAdes: a new genome assembly algorithm and its applications to single-cell sequencing. J. Comput. Biol., 19, 455–477.

30. Li, H., Handsaker, B., Wysoker, A., Fennell, T., Ruan, J., Homer, N., Marth, G., Abecasis, G., Durbin, R. and 1000 Genome Project Data Processing Subgroup (2009) The sequence alignment/map format and SAMtools. Bioinformatics, 25, 2078–2079.

31. Seemann, T. (2014) Prokka: rapid prokaryotic genome annotation. Bioinformatics, 30, 2068–2069.

32. McClure, R., Balasubramanian, D., Sun, Y., Bobrovskyy, M., Sumby, P., Genco, C.A., Vanderpool, C.K. and Tjaden, B. (2013) Computational analysis of bacterial RNA-Seq data. Nucleic Acids Res., 41, e140.

33. Burghout, P., Zomer, A., van der Gaast-de Jongh, C.E., Janssen-Megens,E.M., Françoijs,K.-J., Stunnenberg, H.G. and Hermans, P.W.M. (2013) Streptococcus pneumoniae folate biosynthesis responds to environmental CO2 levels. J. Bacteriol., 195, 1573–1582.

34. Goecks, J., Nekrutenko, A., Taylor, J. and Galaxy Team (2010) Galaxy: a comprehensive approach for supporting accessible, reproducible, and transparent computational research in the life sciences. Genome Biol., 11, R86.

35. Langmead, B. and Salzberg, S.L. (2012) Fast gapped-read alignment with Bowtie 2. Nat. Methods, 9, 357–359.

36. Robinson, J.T., Thorvaldsdóttir,H., Winckler, W., Guttman, M., Lander, E.S., Getz, G. and Mesirov, J.P. (2011) Integrative genomics viewer. Nat. Biotechnol., 29, 24–26.

37. Baldi, P. and Long, A.D. (2001) A Bayesian framework for the analysis of microarray expression data: regularized t-test and statistical inferences of gene changes. Bioinformatics, 17, 509–519.

38. Pritchard, J.R., Chao, M.C., Abel, S., Davis, B.M., Baranowski, C., Zhang, Y.J., Rubin, E.J. and Waldor, M.K. (2014) ARTIST: high-resolution genome-wide assessment of fitness using transposon-insertion sequencing. PLoS Genet., 10, e1004782.

39. Prajsnar, T.K., Cunliffe, V.T., Foster, S.J. and Renshaw, S.A. (2008) A novel vertebrate model of Staphylococcus aureus infection reveals phagocyte-dependent resistance of zebrafish to non-host specialized pathogens. Cell. Microbiol., 10, 2312–2325.

40. Laarman, A.J., Bardoel, B.W., Ruyken, M., Fernie, J., Milder, F.J., van Strijp, J.A.G. and Rooijakkers, S.H.M. (2012) Pseudomonas aeruginosa alkaline protease blocks complement activation via the classical and lectin pathways. J. Immunol., 188, 386–393.

41. Berends, E.T.M., Dekkers, J.F., Nijland, R., Kuipers, A., Soppe, J.A., van Strijp, J.A.G. and Rooijakkers, S.H.M. (2013) Distinct localization of the complement C5b-9 complex on Gram-positive bacteria. Cell. Microbiol., 15, 1955–1968.

42. Pence, M.A., Rooijakkers, S.H.M., Cogen, A.L., Cole, J.N., Hollands, A., Gallo, R.L. and Nizet, V. (2010) Streptococcal inhibitor of complement promotes innate immune resistance phenotypes of invasive M1T1 Group A Streptococcus. J. Innate Immun., 2, 587–595.

43. Prajsnar, T.K., Renshaw, S.A., Ogryzko, N.V., Foster, S.J., Serror, P. and Mesnage, S. (2013) Zebrafish as a novel vertebrate model to dissect enterococcal pathogenesis. Infect. Immun., 81, 4271–4279.

44. Samant, S., Lee, H., Ghassemi, M., Chen, J., Cook, J.L., Mankin, A.S. and Neyfakh, A.A. (2008) Nucleotide biosynthesis is critical for growth of bacteria in human blood. PLoS Pathog., 4, e37.

45. Le Breton, Y., Mistry, P., Valdes, K.M., Quigley, J., Kumar, N., Tettelin, H. and McIver, K.S. (2013) Genome-wide identification of genes required for fitness of group A Streptococcus in human blood. Infect. Immun., 81, 862–875.

46. Jenkins, A., Cote, C., Twenhafel, N., Merkel, T., Bozue, J. and Welkos, S. (2011) Role of purine biosynthesis in Bacillus anthracis pathogenesis and virulence. Infect. Immun., 79, 153–166.

47. Polissi, A., Pontiggia, A., Feger, G., Altieri, M., Mottl, H., Ferrari, L. and Simon, D. (1998) Large-scale identification of virulence genes from Streptococcus pneumoniae. Infect. Immun., 66, 5620–5629.

48. Wang, N., Ozer, E.A., Mandel, M.J. and Hauser, A.R. (2014) Genome-wide identification of Acinetobacter baumannii genes necessary for persistence in the lung. mBio, 5, e01163–01114.

49. Fuller, T.E., Kennedy, M.J. and Lowery, D.E. (2000) Identification of Pasteurella multocida virulence genes in a septicemic mouse model using signature-tagged mutagenesis. Microb. Pathog., 29, 25–38.

50. Schwager, S., Agnoli, K., Kothe, M., Feldmann, F., Givskov, M., Carlier, A. and Eberl, L. (2013) Identification of Burkholderia cenocepacia strain H111 virulence factors using nonmammalian infection hosts. Infect. Immun., 81, 143–153.

51. Chaudhuri, R.R., Peters, S.E., Pleasance, S.J., Northen, H., Willers, C., Paterson, G.K., Cone, D.B., Allen, A.G., Owen, P.J., Shalom, G., et al. (2009) Comprehensive identification of Salmonella enterica serovar Typhimurium genes required for infection of BALB/c mice. PLoS Pathog., 5, e1000529.

52. Mulhbacher, J., Brouillette, E., Allard, M., Fortier, L.-C., Malouin, F. and Lafontaine, D.A. (2010) Novel riboswitch ligand analogs as selective inhibitors of guanine-related metabolic pathways. PLoS Pathog., 6, e1000865.

53. Mainardi, J.-L., Fourgeaud, M., Hugonnet, J.-E., Dubost, L., Brouard, J.-P., Ouazzani, J., Rice, L.B., Gutmann, L. and Arthur, M. (2005) A novel peptidoglycan cross-linking enzyme for a beta-lactam-resistant transpeptidation pathway. J. Biol. Chem., 280, 38146–38152.

54. Theilacker, C., Sava, I., Sanchez-Carballo, P., Bao, Y., Kropec, A., Grohmann, E., Holst, O. and Huebner, J. (2011) Deletion of the glycosyltransferase bgsB of Enterococcus faecalis leads to a complete loss of glycolipids from the cell membrane and to impaired biofilm formation. BMC Microbiol., 11, 67.

55. Davlieva, M., Zhang, W., Arias, C.A. and Shamoo, Y. (2013) Biochemical characterization of cardiolipin synthase mutations associated with daptomycin resistance in enterococci. Antimicrob. Agents Chemother., 57, 289–296.

56. Vollmer, W., Joris, B., Charlier, P. and Foster, S. (2008) Bacterial peptidoglycan (murein) hydrolases. FEMS Microbiol. Rev., 32, 259–286.

57. Mikalsen, T., Pedersen, T., Willems, R., Coque, T.M., Werner, G., Sadowy, E., van Schaik, W., Jensen, L.B., Sundsfjord, A. and Hegstad, K. (2015) Investigating the mobilome in clinically important lineages of Enterococcus faecium and Enterococcus faecalis. BMC Genomics, 16, 282.

58. van Schaik, W., Top, J., Riley, D.R., Boekhorst, J., Vrijenhoek, J.E.P., Schapendonk, C.M.E., Hendrickx, A.P.A., Nijman, I.J., Bonten, M.J.M., Tettelin, H., et al. (2010) Pyrosequencing-based comparative genome analysis of the nosocomial pathogen Enterococcus faecium and identification of a large transferable pathogenicity island. BMC Genomics, 11, 239.

59. Matos, R.C., Lapaque, N., Rigottier-Gois,L., Debarbieux, L., Meylheuc, T., Gonzalez-Zorn,B., Repoila, F., Lopes, M., de F. and Serror, P. (2013) Enterococcus faecalis prophage dynamics and contributions to pathogenic traits. PLoS Genet., 9, e1003539.

60. Duerkop, B.A., Clements, C.V., Rollins, D., Rodrigues, J.L.M. and Hooper, L.V. (2012) A composite bacteriophage alters colonization by an intestinal commensal bacterium. Proc. Natl. Acad. Sci., 109, 17621–17626.

